# Mechanical gating of the auditory transduction channel TMC1 involves the fourth and sixth transmembrane helices

**DOI:** 10.1101/2021.12.30.474567

**Authors:** Nurunisa Akyuz, K. Domenica Karavitaki, Bifeng Pan, Panos I. Tamvakologos, Kelly P. Brock, Yaqiao Li, Debora S. Marks, David P. Corey

**Affiliations:** Department of Neurobiology, Harvard Medical School, Boston, MA 02115, USA; Department of Systems Biology, Harvard Medical School, Boston, MA 02115, USA

## Abstract

The trans<u>m</u>embrane <u>c</u>hannel-like (TMC) 1 and 2 proteins play a central role in auditory transduction, forming ion channels that convert sound into electrical signals. However, the molecular mechanism of their gating remains unknown. Here, using predicted structural models as a guide, we probed the effects of twelve mutations on the mechanical gating of the transduction currents in native hair cells of *Tmc1/2*-null mice expressing virally introduced TMC1 variants. Whole-cell electrophysiological recordings revealed that mutations within the pore-lining transmembrane (TM) helices 4 and 6 modified gating, reducing the force sensitivity or shifting the open probability of the channels, or both. For some of the mutants, these changes were accompanied by a change in single-channel conductance. Our observations are in line with a model wherein conformational changes in the TM4 and TM6 helices are involved in the mechanical gating of the transduction channel.

## Introduction

Our sense of hearing relies on the conversion of sound-induced mechanical stimuli into neural signals by cochlear hair cells of the inner ear. This process hinges on the ability of sensory transduction channels in hair cells to respond to mechanical stimuli by opening and closing, thereby generating fluctuating receptor currents^1^. Despite decades of research, it was difficult to determine the molecular identity of the transduction channel. A number of proteins including trans<u>m</u>embrane <u>c</u>hannel-like 1 and 2 (TMC1 and TMC2), <u>li</u>poma <u>H</u>MGIC fusion <u>p</u>artner-like 5 (LHFPL5), transmembrane inner <u>e</u>ar <u>e</u>xpressed protein (TMIE) and <u>c</u>alcium and integrin <u>b</u>inding <u>p</u>rotein (CIB2) are now known to be essential for mechanotransduction ^2, 3, 4, 5^ but their specific roles were not clear.

TMCs can assemble as dimers in solution and are structurally related to the TMEM16 and TMEM63/OSCA proteins^6, 7, 8^. Each subunit of the dimer is thought to contain ten transmembrane (TM) helices (**Supplementary Fig. 1A-B**). Parts of the N-terminus (residues 100-130 in mouse TMC1) as well as the intracellular loop between TM helices 2-3 (residues 300-350) have been implicated in binding to the auxiliary channel subunit CIB2 ^3, 9^. Currently, we do not know the mode of dimerization; however the models based on X-ray and cryo-EM structures of TMEM16 proteins^6, 8^ suggest that the subunit interface is near TM10 and may be at least partly filled with lipid as has been observed in TMEM16 and OSCA protein families (**Supplementary Fig. 1B**)^10^ or perhaps by auxiliary proteins^11^.

Experiments with spontaneous and engineered mutations of the TMC1 protein have now indicated that TMC1 (and likely TMC2) is a pore-forming protein of the mechanotransduction complex^5, 6^. Each subunit of the dimer includes a separate ion-conducting pore, which is a large open cavity that is surrounded by TM helices 4-7 (**Supplementary Fig. 1B**) ^6, 8, 12^. Channel activity of TMC homologs expressed in heterologous cell lines^13^ is also consistent with the idea that TMC1 and TMC2 constitute most or all of the transduction channel pore.

These experiments did not, however, shed light on gating—on the mechanically evoked conformational changes that open and close the pore. Conformational states associated with conduction have been suggested for the related ion channels and lipid scramblases of the TMEM16 family. Specifically, in TMEM16 chloride channels and lipid scramblases, Ca^2+^-activated opening is thought to involve an initial rearrangement of the ‘gating helix’ TM6^14, 15^. Channel opening further requires disruption of interactions between TM4 and TM6^16, 17^.

Molecular dynamic simulations of TMC1 structural models based on TMEM16 proteins suggest that the TMC1 pore can adopt two distinct conformations. In one conformation, TM4 are about 4 Å further from TM6 than in the other, and in simulations of K^+^ ion passage through the pore, the more open conformation conducted ions about three times faster^12^. These studies provide starting points for probing the gating of TMC1 physiologically.

In this study, for the first time, we probe the relationship between the structure of TMC1 and the mechanical activation of the hair-cell transduction channel. We used structure-prediction algorithms to further explore potential conformations of TMC1. Based on these, we designed mutations in TM4 and TM6 that might affect gating. We used AAV vectors to express, in hair cells of *Tmc1/2* double-knockout (DKO) mice, TMC1 channels bearing specific mutations and we recorded mechanotransduction currents from cochlear hair cells. We examined the effects of a dozen mutations on mechanical gating of the transduction currents in mouse hair cells in vivo (**Supplementary Table 1)**. Shifts in the mechanical activation curves suggested that many of these mutations changed the relative energy of the open state. Furthermore, some of the mutations reduced the slope of the activation curve, suggesting that these mutations render the channel less sensitive to mechanical stimulus. In some cases, these gating changes were accompanied by a decrease in single-channel conductance, measured with nonstationary noise analysis, consistent with TM4 and TM6 also contributing to the permeation pathway^6^. Together, these observations provide evidence for a model of transduction wherein conformational changes involving TM4 and TM6 of TMC1 are critical for channel opening induced by hair bundle deflection.

## Results

### Predicted TMC1 Structures

To identify residues that may influence gating, we began with our previous homology model for TMC1^6^. We then generated more refined models using the deep learning-based modeling algorithms, transform-restrained (tr) Rosetta^18^ and RoseTTAFold ^19^ (see Methods, **Fig. 1A**). These methods rely on patterns in protein sequences as well as on amino-acid interactions suggested by co-evolution, and utilize energy minimization techniques^20^ to predict protein structures. Without relying on structures of related proteins, both trRosetta and RoseTTAFold predict an overall ‘TMEM16-like’ fold for TMC1 that is very similar to that previously predicted from homology to TMEM16s^6^ (**Fig. 1A**).

**Figure 1.**
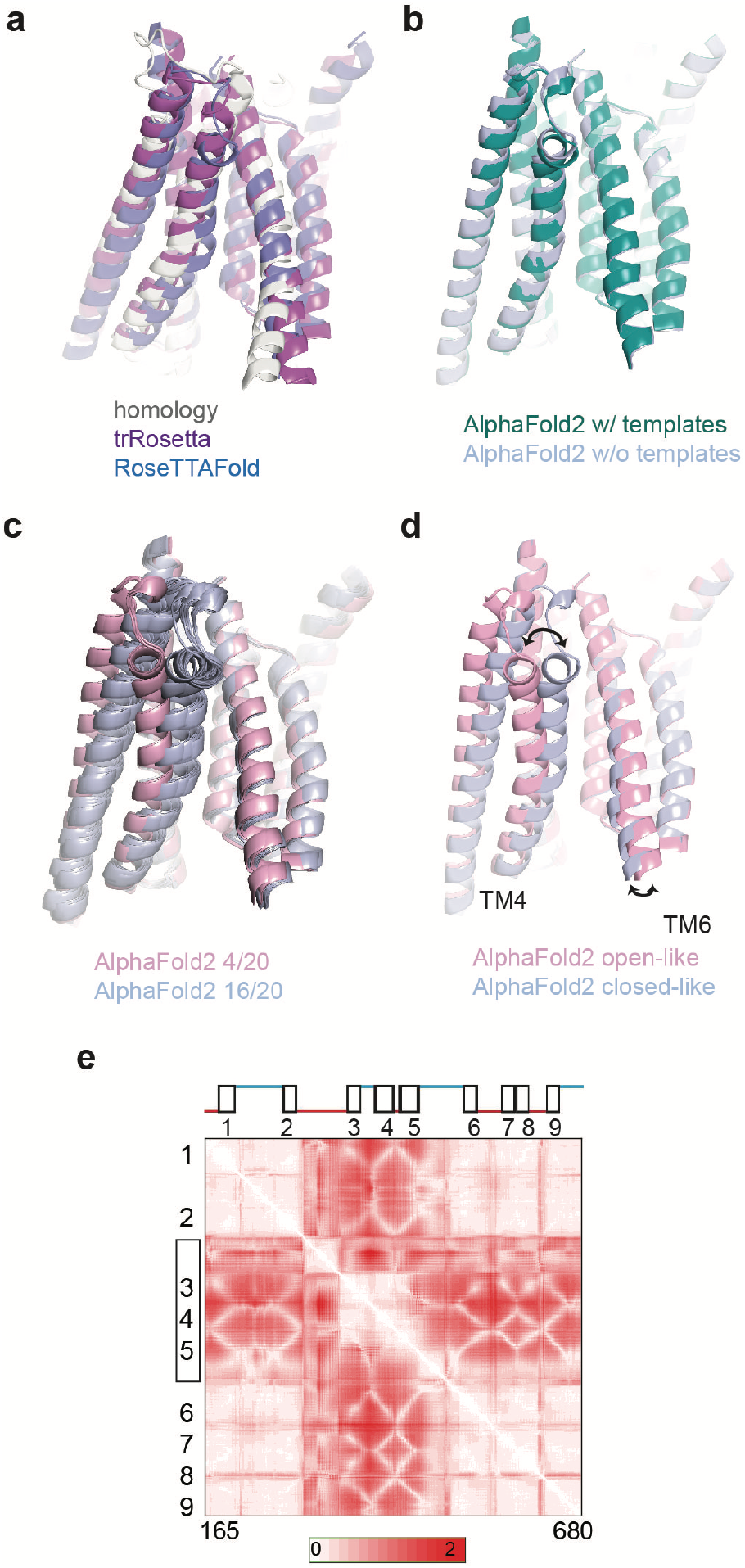
Similarity in TMC1 structural predictions. **a** Structural models for mouse TMC1. Shown in foreground are TMs 3,4,6 and 8, viewed from within the membrane plane. Gray represents an iTasser model based on homology to TMEM16 channels. Purple shows a prediction by trRosetta, not based on other structures. Blue is a predicted model by RoseTTAFold, also not based on known structural templates. The three models are nearly identical. **b** Structural models for TMC1 from AlphaFold2. Green shows a model based on templates such as TMEM16A. Blue is the ab initio model not based on known structures. In the pore region they are the same within 1-2 Å. **c** Two models from AlphaFold2, generated with multiple seed parameters. In 20 model iterations, 16 grouped in one conformation (blue; like that in panel B) and 4 grouped in a different conformation (pink). **d** Representative models from the two groups of conformations in panel C. The conduction pathway for ions is between TM4 and TM6. In the more open conformation (pink), TM4 and TM5 are more distant from TM6 by about 5 Å at the constricted region of the pore. **e** Standard-deviation (SD) maps for all residue-residue (R-R) distances for residues 165-680. For each residue pair, more intense red represents a larger change in distance between the two conformations. TM helices are numbered. The N-terminus, C-terminus and TM10 were not included in the analysis.

We then performed predictions of the TMC1 structure using the AlphaFold2 program developed by DeepMind^21, 22^ (**Fig. 1B**). The predictions with or without utilizing a structural template were nearly identical and were very similar to previous models, especially in regard to the overall fold of the pore region (**Fig. 1B**).

These models converge on a pore region in each subunit of the dimer that is formed by TM4-7 and exhibits an overall negative electrostatic potential, which is consistent with the cation selectivity of the channel^23, 24, 25^. Several conserved residues within TM6-7 of mTMC1 contribute to this negative surface charge, including E514, E520, D528, D540, D557, E559, D569, E567. Mutations of some of these residues, including D528 and D569, have been linked to hearing loss ^26, 27^.

Running the AlphaFold2 network multiple times using different seeds, we were able to increase the diversity of the predicted structural states (**Fig. 1C**). We generated 20 models (4 random seeds and 5 models from each), and then aligned and clustered these structures using Chimera^2^. By calculating the root-mean-square deviation (RMSD) values for every pair-wise comparison between the 20 models, we found the structures clustered into two groups (16 structures in one group and 4 in the other). In the first group, TM3 and TM4 were close to TM6 (‘closed-like’; **Fig. 1D**). In the second, TM3 and TM4 maintained a close association with each other but were further from TM6 (‘open-like’). The distance between the C-α atoms of residues 409 (on TM4) and 531 (on TM6) within the pore region is larger by ∼5 Å in the open-like structures. The HOLE software^28^ consistently showed a water-filled pore conformation with a large conduction pathway for the open-like but not closed-like predicted structures (**Supplementary Fig. 2A**). The narrowest part of the pathway in this model occurs where N404 in TM4 approaches the positively charged R523 and the hydrophobic L524 in TM6. A thin cross-section of the protein parallel to the membrane near N404 and L524 displays the opening clearly (**Supplementary Fig. 2B**). Notably, the change between the predicted states seems to be driven primarily by a ∼10° tilting and translation of TM4 together with TM3, but also by smaller movements in TM6 (**Fig. 1D**). Interestingly, even the open-like structures displayed a pore diameter of ∼ 6Å in the narrowest dimension, which is smaller than that estimated for this channel based on the sizes of the permeating molecules ^8, 29^.

We then generated 160 structures (32 random but distinct seeds and 5 models each) and mapped residue-residue (R-R) distances to compare the AlphaFold2 generated structures^30^. These maps showed that across the 160 structures there was a large variability in the position of the TM3-5 helices with respect to the rest of the protein (TMs 1,2,6-9; **Fig. 1E**). Most of the intracellular TM2-TM3 loop shows variability with the rest of the protein, as does the first half of the TM5-6 extracellular loop. However, these two loops also show variability with TM3-5. If the differences between the open-like and closed-like structures predicted by AlphaFold2 in fact represent a gating transition, there may be three distinct domain movements associated with gating.

We also generated contact maps by EVcouplings software^31^ (see Methods) and mapped them onto the open-like and closed-like structure predictions. Evolutionarily coupled residue pairs (ECs) are found in close proximity in the models, as expected (**Supplementary Fig. 2C)**. However, there are notably more contacts (not just ECs) in the closed-like state (**Supplementary Fig. 2C)**.

Overall, the models generated by AlphaFold2 lead us to hypothesize that in TMC1, as in the structurally similar proteins of the TMEM16 and TMEM63/OSCA families, the pore helices TM4 and TM6 separate during gating^15, 32, 33^.

### Activation curves of virally expressed TMC1 in cochlear hair cells of Tmc1/2-DKO animals

If the pore helices TM4 and TM6 undergo a global rearrangement during gating, then mutations of critical residues within these helices may change the activation of the channel. We therefore designed and assessed the effects of mutations in this region of TMC1 (**Fig. 2A-B, Supplementary Table 1**). We generated AAV9-PHP.B viral vectors carrying a sequence encoding either wild-type (WT) or mutated TMC1 (**Fig. 2C**). Viral vectors were individually injected into the inner ears of *Tmc1/2-*DKO animals at postnatal day (P)1, using the round window approach (see Methods; **Fig. 2C**). Cochlear dissections were performed at P4–P6. The tissue was placed in culture for an additional 3–7 days (**Fig. 2C**, bottom panel). The mid-basal section was used for electrophysiology and the mid-apical section was used to assess viral transduction efficiency and functional rescue using an FM1-43 dye loading assay (see Methods). At this stage of cochlear development, *Tmc2* is mostly down-regulated, but we used *Tmc1/2-*DKO animals to avoid the possibility of any contribution of TMC2 to transduction currents.

**Figure 2.**
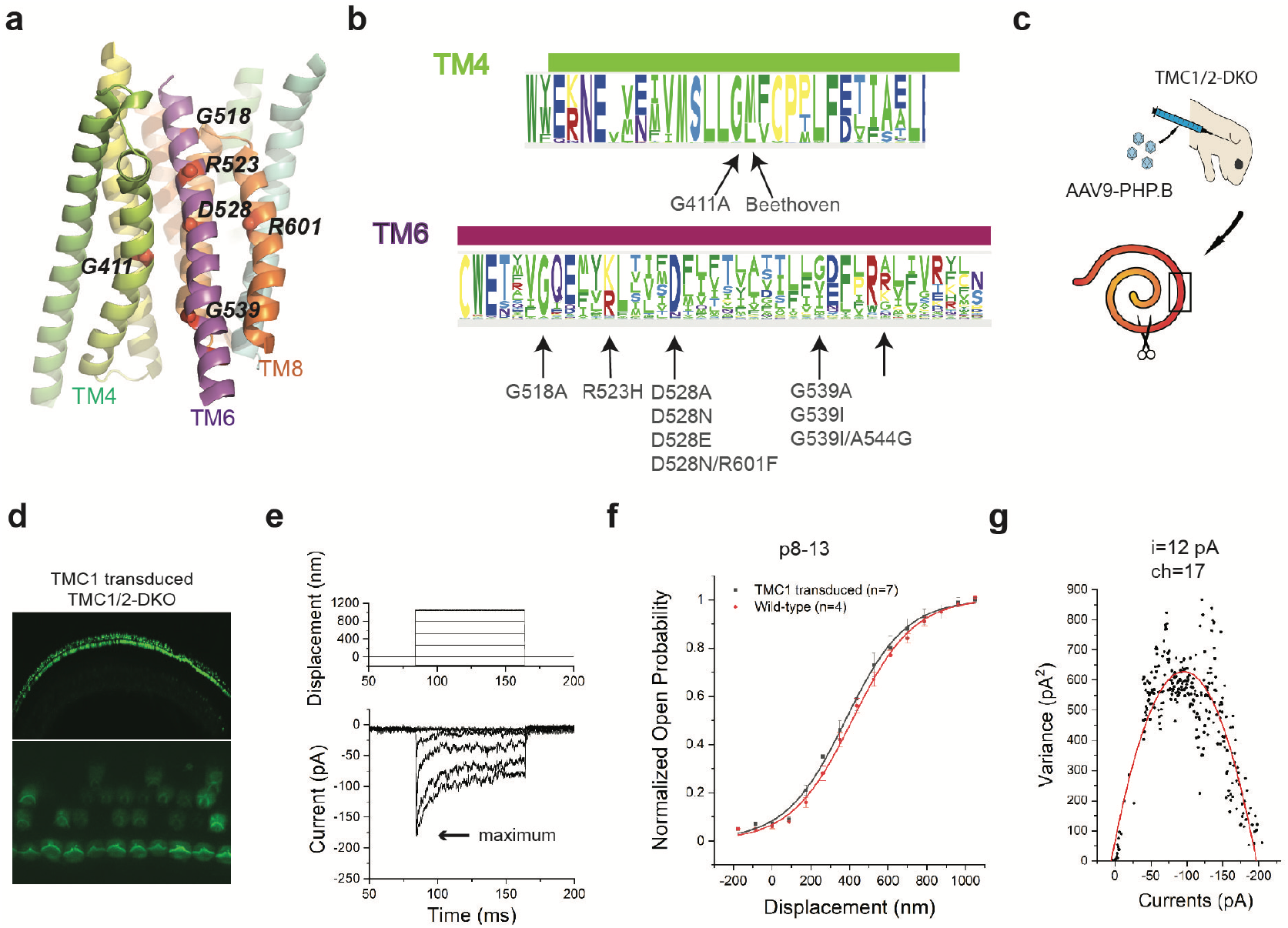
Structure-based mutations of TMC1 and functional assays. **a** Locations of the mutation sites in a view of the pore region from the plane of the membrane. **b** A sequence logo generated from multiple sequence alignments for TM4 and TM6 regions of TMCs; mutations in this study indicated. **c** Cartoon demonstrating AAV injection of neonatal *Tmc1/2* double-knockout (DKO) animals (top), and separation of the dissected cochlear explants into two sections (bottom). The scissors indicate where the tissue was split. The black box represents the mid-base region, used for electrophysiological recordings of transduction currents. **d** FM1-43 loading of cochlear HCs from *Tmc1/2-*DKO mice expressing virally expressed WT TMC1 channels. The bottom panel is a magnified view at the plane of the bundles. **e** Representative transduction currents in response to -175 to 1050 nm bundle deflections (top panel) recorded from DKO IHCs expressing virally encoded WT-TMC1. **f** Activation curves. Normalized open probability curves are shown as a function of stimulus displacement (nm), in WT-mice and *Tmc1/2-*DKO mice expressing WT-TMC1. The curves are fitted with a Boltzmann equation. **g** Single channel current estimate (i=12 pA) and number of channels (ch=17) based on non-stationary noise analysis from a hair cell expressing viral WT-TMC1. Variance is plotted as a function of current and fitted with a parabolic equation.

Consistent with previous findings, we observed that cochlear hair cells of *Tmc1/2-*DKO mice do not take up FM1-43, indicating that they lack functional transduction channels ^34, 35^. In contrast, *Tmc1/2-*DKO hair cells transduced with AAV9-PHP.B-*Tmc1* encoding WT-TMC1 showed FM1-43 uptake, indicating rescue of transduction activity (**Fig. 2D**). Specifically, the FM1-43 assay indicated that ∼ 90% of inner hair cells (IHCs) expressed functional TMC1 channels. This is in line with recent studies demonstrating the use of AAV9-PHP.B vectors with CMV promoters as highly efficient for driving expression in hair cells ^36, 37^.

Following FM1-43 screening, electrophysiological recordings were performed using a stiff glass probe to stimulate hair bundles and whole-cell patch clamping to record currents, as previously described ^6^. First, we recorded currents from IHCs of wild-type mice at P8-P13. Maximum current amplitudes of 660±80 pA (n=4) were obtained. In contrast, DKO hair cells expressing virally expressed WT TMC1 channels yielded receptor currents with maximal amplitudes of 202±33 pA, n=7 (**Fig. 2E**). The amplitude and variability in the whole cell currents were consistent with previous reports ^6, 36^.

To obtain activation curves for TMC1 channels, we measured IHC currents evoked by a series of 15 bundle step deflections ranging from -175 to 1050 nm (see Methods). Since the shape and slope of the activation curves reflect properties of individual channels, we normalized the currents for each cell to allow for comparison to other cells (**Fig. 2F**).

We fitted the normalized activation curves of the channels with a Boltzmann equation:

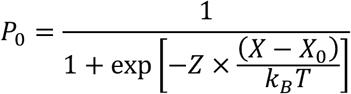

Here, Z represents the apparent mechanical sensitivity of the channel; it determines the slope of the activation curve ^38^. *X*_0_ represents the bundle deflection at which the channel open probability is 0.5. *k*_B_*T* is thermal energy (4.1 pN nm at room temperature). The normalized activation curves that we obtained from IHCs of TMC1-injected *Tmc1/2-*DKO mice were consistent with previous observations using the same stimulation technique ^5, 6^ (**Fig. 2F)**. The fits of the activation curves of virally transduced WT-TMC1 in IHCs of *Tmc1/2-*DKO animals yielded Z values of 0.027 ± 0.003 pN and X_0_ values of 380± 32 nm.

Using nonstationary noise analysis of whole cell currents^39, 40^ we estimated single channel currents as 12.5 ± 2.5 pA (**Fig. 2G**), which are also in line with our previous measurements ^6^.

### Mutation of a glycine residue on TM4 alters the activation curve

We first targeted glycines within TM4 and TM6 because glycines within transmembrane alpha helices of ion channels tend to act as hinge points for a gating conformational change ^41^. There is one glycine within TM4 (G411)^26^ which is highly conserved across TMCs (**Fig. 2B, Fig. 3A**); mutations of this residue have been associated with deafness. We mutated this residue to a helix-stabilizing alanine^42^ to assess whether reducing the conformational flexibility of TM4 would affect gating.

**Figure 3.**
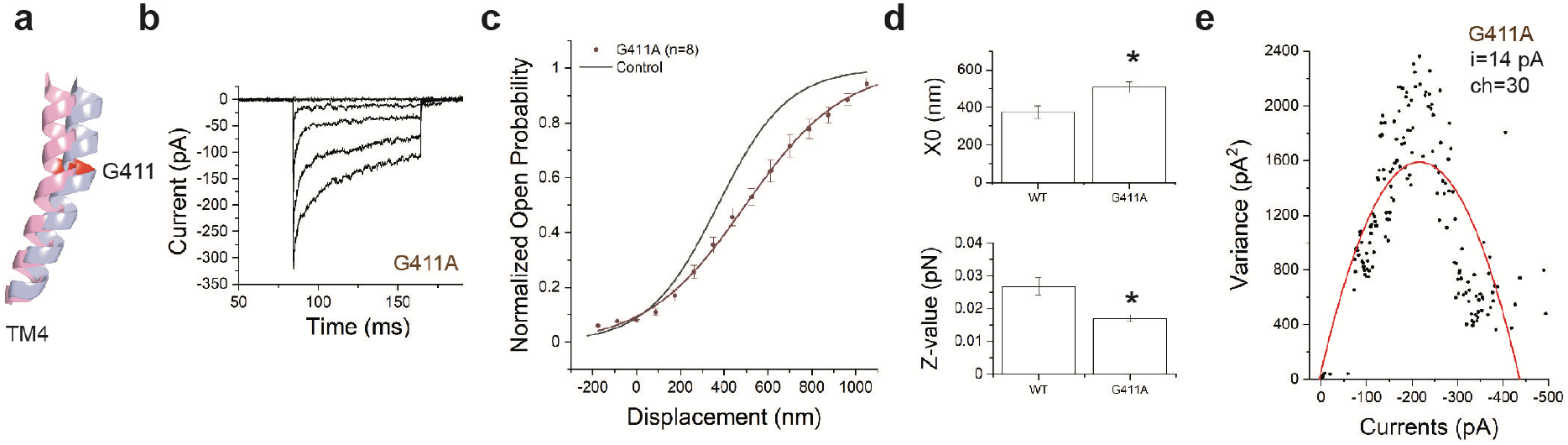
Mutation of G411 on TM4 shifts the activation curve and changes the slope. **a** Representation of TM4 showing G411A mutation site. Pink and blue represent the open- and closed-like configurations, respectively. **b** Representative transduction currents in response to -175 to 1050 nm deflections from IHCs expressing TMC1-G411A. **c** Activation curves from TMC1-G411A (red) and WT-TMC1 (black) expressed in DKO mice. **d** Fitting parameters X_0_ (position of half-activation) and Z (slope factor). Significance in differences in X_0_ (p=0.01) and Z-values (p=0.01) are indicated stars above the bars. **e** Non-stationary noise analysis for a hair cell expressing TMC1-G411A. Conductance did not change significantly.

Recordings from IHCs expressing TMC1 channels bearing the G411A mutation yielded maximum currents of 291 ± 38 pA (n=8), which are comparable to those obtained from WT-TMC1 injected IHCs (**Fig. 3B)**. However, the activation curves of TMC1-G411A were shifted rightward (*X*_0_ = 507 ± 28 nm) compared to WT-TMC1 (*X*_0_=380 ± 32 nm) (**Fig. 3C**). The slopes of the curves, represented by the Z values, were also significantly decreased—from to 0.027 to 0.017 pN. Both these changes were significant with p-values <0.05 (**Fig. 3D)**. The single channel current estimated from noise analysis was 14 ± 3.5 pA (**Fig. 3E**), which is close to the estimate for WT-TMC1^6^. Taken together, these findings are consistent with the idea that the G411A mutation influences the mechanical gating but not the unitary conductance of the channel.

### Mutations of glycine residues on TM6 shift the activation curves

We then turned to the glycine residues in TM6. There are two: G518 is close to the extracellular side of the helix, and G539 is closer to the intracellular side (**Fig. 2A-B, Fig. 4A**). From IHCs expressing TMC1-G518A, we obtained maximum currents of 211± 11 pA (n=8) which are comparable to those from WT-TMC1 (**Fig. 4B, Supplementary Fig. 3A**). TMC1-G518A activation curves were shifted to the right by ∼110 nm (*X*_0_= 493±16 nm) compared to WT-TMC1 (*X*_0_=380 ± 32 nm) (**Fig. 4C-D**). From IHCs expressing TMC1-G539A, we also obtained typical maximum currents of 214 pA ± 38 pA (n=14) and the activation curves were shifted rightward by ∼140 nm (*X*_0_ = 518 ± 30 nm) (**Fig. 4C**). For both TMC1-G518A and TMC1-G539A, the difference in *X*_0_ compared to WT-TMC1 is statistically significant, with p<0.05 (**Fig. 4D)**.

**Figure 4.**
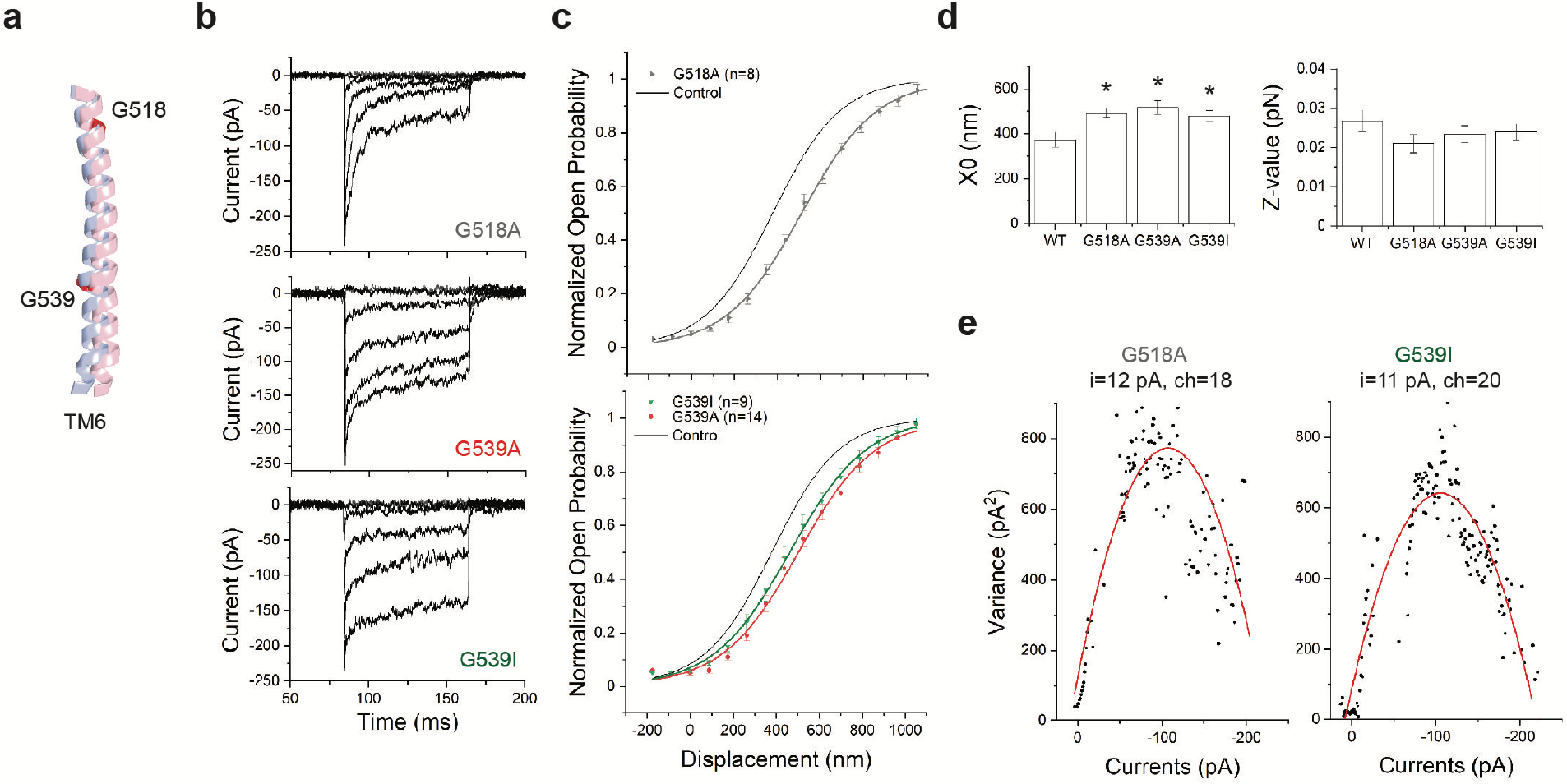
Mutation of G518 or G539 on TM6 shifts the activation curves without changing the slope. **a** Relative position of TM6 in open-like (pink) and closed-like (blue) structural models generated by AlphaFold2, showing G518 and G539. The predicted structures were aligned globally. **b** Representative transduction currents in IHCs expressing TMC1-G518A (top), TMC1-G539A (middle) and TMC1-G539I (bottom). **c** Activation curves from TMC1-G518A (top) and TMC1-G539A and TMC1-G539I (bottom). In both, the control WT-TMC1 activation curve is shown in black. **d** Fitting parameters X_0_ and Z. All three mutations shifted the activation curves (with p-values = 0.03, 0.01, 0.02) without significantly changing slope (with p-values = 0.08, 0.44, 0.13, respectively). **e** Non-stationary noise analysis for single hair cells expressing G518A (left) or G539I (right). Conductance did not significantly change from wild-type.

Importantly, for both these mutations in TM6, the Z values were not significantly different from those for wild-type TMC1 (**Fig. 4D**). A change in *X*_0_ without a change in Z suggests that these mutations in TM6 alter the open probability of the channel without changing the way it is coupled to the mechanical stimulus. Using a simple two-state model (see Methods), these changes in *X*_0_ can be interpreted as an increase in the energy of the open state compared to the closed state of ∼0.5 k_B_T, thus favoring the closed state. Finally, neither TMC1-G518A nor TMC1-G539A showed a significant effect on single channel currents, with 12 ± 2.5 pA for G518A and 14 ± 2.5 pA for G539A (**Fig. 4E**).

The glycine at position 518 is highly conserved: 98% of the TMC sequences in our multiple sequence alignment (MSA) show a glycine at this position. However, only 50% of the sequences have a glycine at position 539. A significant portion of the remaining sequences (41%) have an isoleucine or a valine at this position (**Supplementary Table 2**). These sequences include the closely related mouse TMC3, in which the amino acid corresponding to G539 is an isoleucine (**Supplementary Fig. 3B**). We found that the effect of a G539I mutation in TMC1 was similar to that of G539A (**Fig. 4B-D**): IHCs expressing TMC1-G539I showed normal transduction current amplitude of 190 pA ± 14 pA (n=9) and single channel currents (14 ± 3 pA) but a ∼100 nm rightward shift in the activation curves (*X*_0_ = 478 ± 25 nm) compared to WT-TMC1 (**Fig. 4C-D**). Again, no significant change was observed in the Z value, implying that this mutation, similar to TMC1-G539A, does not affect the mechanical sensitivity of the channel **(Fig. 4C-D)**.

TMC3 has a glycine four residues towards the intracellular side of the protein, equivalent to position 544 in TMC1 (**Fig. 2B, Supplementary Fig. 3B**). We wondered whether a glycine anywhere in this region would provide sufficient flexibility, and so re-introduced a glycine at position 544 in TMC1 lacking the glycine at position 539 (TMC1-G539I/A544G). However, we found that a glycine at position 544 did not substitute well for one at position 539, and the activation curve of TMC1-G539I/A544G was not significantly different from TMC1-G539I alone (**Supplementary Fig. 3C-E**).

We conclude that mutation of either of the glycine residues at positions 518 and G539 on TM6 reduces the resting open probability, as indicated by the rightward shift in *X*_*0*_, but does not contribute to the mechanical sensitivity of the channel as measured by the slope Z, or to ion permeation. This is consistent with the idea that flexibility of TM6 is important for channel opening.

### D528 on TM6 is Critical for Mechanical Sensitivity and Ion Conduction

TM6 contains a central and highly conserved aspartate, D528 (**Fig. 2A-B**). *Tmc1* p.D528N is a recessive deafness mutation in mice^26^. The corresponding residue within the Ca^2+-^activated Cl^-^ channel TMEM16A is a lysine (K645 in mouse TMEM16A), which has been linked to the voltage, Ca^2+^ and anion dependence of the channel activation ^32^. In the structural models for TMC1, D528 faces the channel pore^6, 8, 26^ and significantly contributes to the negative surface charge (**Fig. 5A, Supplementary Fig. 4A**). To investigate whether D528 plays a role in mechanosensitive gating, we made the following mutations: D528A, D528N and D528E.

**Figure 5.**
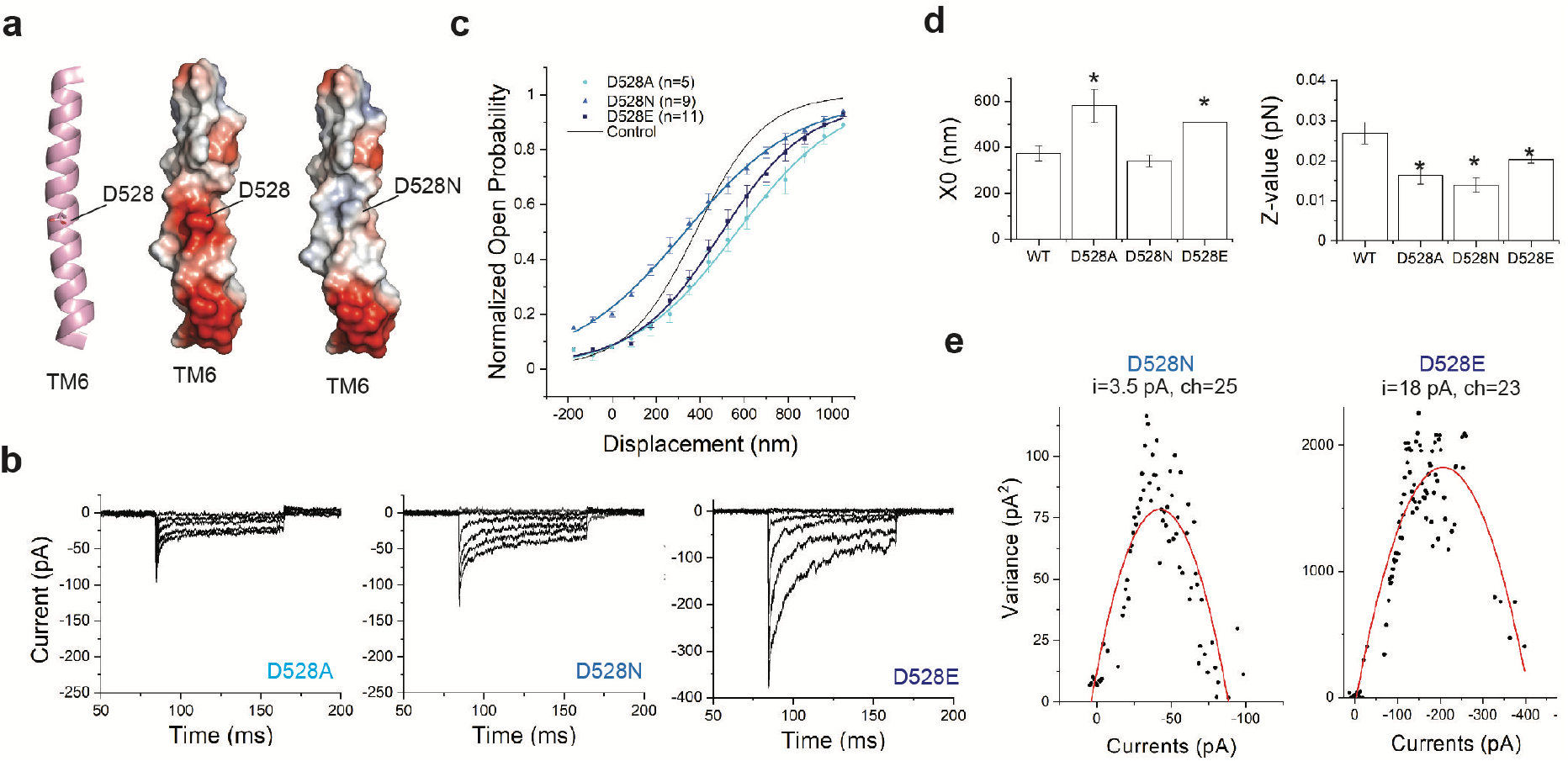
Mutations of D528 shift the activation curve and change the slope, and can change single channel currents. **a** Position of D528 within TM6 of TMC1. TM6 is shown cartoon representation (left panel) with the side-chain of D528 highlighted in stick. The right panels show TM6 in electrostatic surface representation. Electrostatic surface potentials are colored red and blue for negative and positive charges, respectively, and white color represents neutral residues. **b** Transduction currents from TMC1 bearing D528A (left), D528N (middle) or D528E (right) mutations. **c** Activation curves from IHCs expressing TMC1-D528A (cyan), D528N (blue) or D528E (navy). The fit for WT-TMC1 is shown in black as reference. **d** X_0_ and Z for each mutation compared to WT. All three mutations reduced the slope (p-values<0.01), whereas the midpoints of curves for D528A and D528E (p-values= 0.01, 0.01), but not D528N (p-value=0.33), were shifted. **e** Non-stationary noise analysis for TMC1-D528N (left) and TMC1-D528E (right). Preservation of charge in the D528E mutant preserved single-channel conductance.

These mutations are also expected to change the surface electrostatics within the pore cavity (**Supplementary Fig. 4A**). Indeed, we had previously found that IHCs expressing TMC1-D528C show significantly reduced transduction currents (∼30% relative to WT-TMC1)^6^. The currents were further diminished—irreversibly—with application of MTSET reagents (down to ∼2% relative to wild-type), confirming their location within the pore^6^.

Here, we first mutated D528 to an alanine to assess the effect of removing the aspartate side chain. We obtained average maximum currents of 80 ± 15 pA (n=5), which are significantly smaller than that for other mutations we examined (**Fig. 5B, Supplementary Fig. 5A**). Remarkably, we found a 40% reduction in the slope of the TMC1-D528A activation curve compared to WT-TMC1 (**Fig. 5C-D**). In addition, the activation curve was shifted to the right by nearly 190 nm (566 ± 70 nm compared to 380 ± 32 nm). This apparent shift in the half-maximum location can be accounted for by the dramatic change in the slope; the resting open probability was not altered.

We then mutated D528 to an asparagine (D528N), which is similar to an aspartate in size but lacks the negative charge. The average maximum current was near normal at 148±18 pA, n=9 (**Fig. 5B, Supplementary Fig. 5A**). We found that the D528N mutation also reduced the slope of the activation curve compared to WT-TMC1 (**Fig. 5C**,**D**). Reduction in slope in both D528A and D528N suggest that the aspartate side chain at this position contributes to the mechanical sensitivity of the channel. Furthermore, there was a leftward shift of the activation curve (X_0_= 340 nm), indicating an increase in the resting open probability of TMC1-D528N (∼20%) compared to WT-TMC1 (∼6%) (**Fig. 5C**).

As D528 is both negatively charged and situated at a central location within the pore, we reasoned that the loss of charge in these mutants would reduce cation permeation. Consistent with another recent study, we found a striking decrease in TMC1-D528N single channel currents: 3.5 ± 0.5 pA compared to 12.5 ± 2.5 pA for WT-TMC1 (**Fig. 5E**) ^6, 26, 43^. We therefore mutated D528 to glutamate, expecting that substitution with a negatively charged residue may have milder effects. IHCs expressing TMC1-D528E showed maximum average transduction currents of 332 pA ± 40 pA (n=11). The large single channel current for this mutant (17± 2.1 pA) suggests that the negative charge of the amino acid at this position is indeed critical for attracting cations to the pore and so contributes to high conductance (**Fig. 5E)**. However, the charge is not the sole determinant of D528’s contribution to channel activation: We found that the TMC1-D528E mutant had a lower slope than WT-TMC1 (**Fig. 5B-D**), suggesting that the size of the side chain of D528 matters for gating and that the role of D528 in gating is more than providing a negative charge. The fact that there is no glutamate at this position in a significant fraction of all the TMC sequences in the multiple sequence alignment is also consistent with this observation (**Supplementary Table 3**). Whether the amino acid size is changed by one carbon (D-to-E) or the size kept but the charge removed (D-to-N), the mechanical sensitivity is affected.

It is possible that the observed effects of D528 mutations may be indirect through a reduction in Ca^2+^ permeation and its effect on the resting open probability. However, a previously characterized deafness mutation, M412K, also leads to a reduction in Ca^2+^ permeation but no change in the mechanical sensitivity of the channel as represented by the slope of the activation curves ^5, 6^, so it is more likely that D528 is directly involved in channel gating.

### Mutations of positively charged residues near D528 do not significantly alter TMC1’s mechanical sensitivity

A positively charged residue near D528 is R601 on TM8. Evolutionary coupling analysis revealed that D528 on TM6 strongly coevolves with R601^6^. Indeed this is the second strongest coupling out of all possible analyzed pairs. Within a multiple sequence alignment of ∼5000 TMC sequences, 90% of sequences have an aspartate at the position equivalent to 528. When it is an aspartate, the amino acid at the position equivalent to 601 is positively charged—either an arginine or a lysine. When 528 is not an aspartate (11%), 601 loses charge and is either a phenylalanine (F) or less frequently an isoleucine (I)—amino acids with hydrophobic chains (**Supplementary Table 3**).

The proximity of D528 on TM6 to R601 on TM8 in the structural models generated by AlphaFold2 suggest that they can form a salt bridge (**Fig. 6A, Supplementary Fig. 4B**). Similarly, in the structurally related hyperosmolality-gated OSCA calcium channels, interactions among TM6, TM7 and TM8 have been implicated in the gating of the channels^44^. In the calcium-gated TMEM16A chloride channels, calcium ions bind among acidic residues in TM6, TM7 and TM8 to activate the channels^14^. To assess the influence of the arginine at 601 on TMC1 gating, we mutated R601 in WT-TMC1 as well as in TMC1-D528N. TMC1-R601A yielded very low whole-cell currents (23 ± 8 pA, n=5), while TMC1-D528N/R601F showed currents (166 ± 40 pA, n=5) comparable to WT-TMC1 (**Fig. 6B, Supplementary Fig. 5A**). In both WT-TMC1 and TMC1-D528N, mutation of R601 shifted the activation curves rightward, apparently stabilizing the closed state (R601A: *X*_*0*_=580±25 nm; D528N/R601F: *X*_*0*_=510±46 nm) (**Fig. 6C-D**). The shift in *X*_0_ is statistically significant for both (**Fig. 6D**). The Z values, representing mechanical sensitivity of the channel, were not affected by these arginine mutations (**Fig. 6D**).

**Figure 6:**
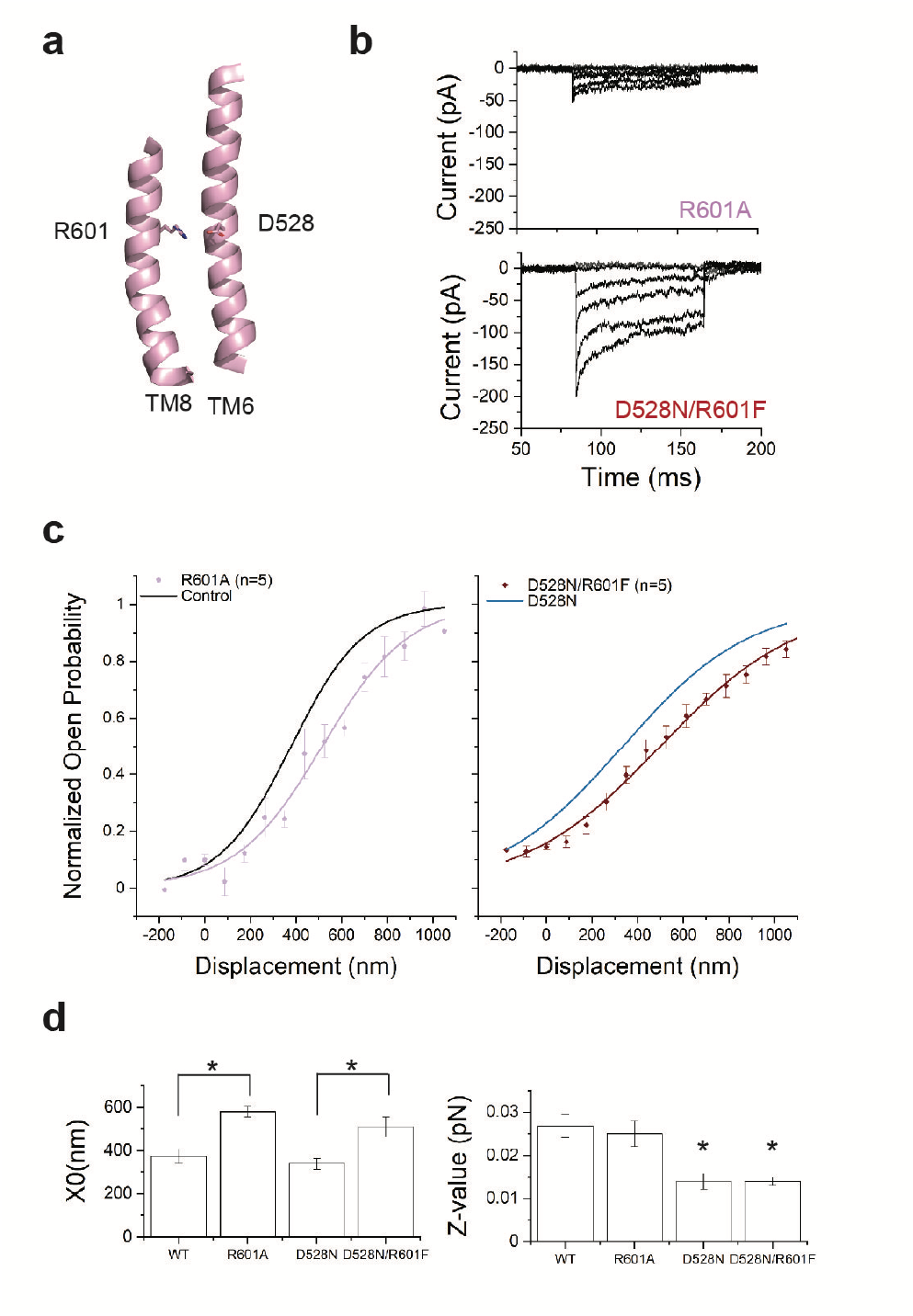
Mutation of R601 shifts the activation curve without changing slope. **a** Representation of TM8 showing R601. **b** Transduction currents from TMC1-R601A (top) and TMC1-D528N/R601F mutation (bottom). **c-d** Activation curves from IHCs expressing TMC1-R601A (lilac) or TMC1-D528N/R601F (crimson). The fits for wild-type TMC1 (C, black) and TMC1-D528N (D, blue) are shown as reference. **d** X_0_ and Z values. R601A shifts the curve (p-value<0.01) from WT without significantly changing slope (p-value=0.1), and shifts the curve from D528N (p-value<0.01) without further changing slope (p-value=0.3).

Thus removing a positive charge from TM8 that is likely involved in a salt bridge to TM6 changes the instrinsic energy difference between the open and closed states, without changing the efficiency with which the mechanical stimulus is coupled to channel gating. This implies further that the dramatic change in the mechanical sensitivity of the channel following mutation of D528, as measured by the slope, cannot be explained by electrostatic interaction between D528 and R601 but is caused by interaction between D528 and other residues.

## Discussion

We previously used cysteine mutagenesis and cysteine modifying reagents to implicate TMC1 transmembrane domains 4-7 in the ion permeation pathway opened by mechanical stimulation^6^. Other studies involving mutations in these domains, including M412K ^5, 45, 46^, D569N ^27^, T416K, W554L and D528N ^27^, are consistent with this implication, as are detailed molecular dynamic simulations of ion permeation using predicted TMC1 structures ^10^. The opening itself, however, is less well understood.

In this study, we used viral gene delivery of mutant TMC1 to TMC1/2-null hair cells to study channel gating. To guide mutations, we first turned to machine-learning based, template-free structure prediction programs, including AlphaFold2^21^, to refine our previous homology model^6^. The newer, ab initio models predicted a fold for TMC1 (**Fig. 1A-D**) very similar to that based on the known structures of the related TMEM16 channels ^6,9^, adding confidence in the models. Running AlphaFold2 multiple times with different starting conditions suggested structural diversity, with striking differences in the pore region of TMC1. These predicted conformations suggest that TM3 and TM4 together move away from TM6 to open the channel (**Supplementary Movie** 1). Similarly, distancing of TM4 and TM6 enhances permeation in molecular dynamics simulations of the homology models^12^.

For TMC1, the idea that conformational changes in TM4 and TM6 might bring their more extracellular portions in close contact and close off the permeation pathway is consistent with a gate for the hair-cell transduction channels that is located outside the binding site for charged blockers ^47^. Previous experiments demonstrated that M412 and D569 sites, which face the more intracellular part of the pore, are accessible from the extracellular solution only when the transduction channels are open but not when they are closed ^6^; they are also consistent with a gate near the outer part of the pore (**Supplementary Fig. 4A**).

Interestingly, the open-like conformations predicted by AlphaFold2 show a pore that is, in its narrowest dimension, not larger than about 6 Å in diameter, which is similar to what has been observed in recent molecular dynamics simulation studies^12^. Careful physiology has suggested a larger open pore diameter, close to 12 Å ^8, 29^. It may be that the pore can expand further than suggested in these models, or it may be that large permeant ions can extend partially into the lipid as they pass through the pore ^6^.

Based on the predicted structures, we prepared a total of twelve mutations at seven sites in TMC1. We found that mutations of a set of glycines within the TM6 helix led to rightward shifts in the activation curve without a change in slope. These glycine mutations lower the resting open probability of the channel, suggesting that they reduce the energy of the closed state without changing the way that force produced by tip link tension is conveyed to the gate.

On the other hand, mutations of the pore-lining residues G411 on TM4 and D528 on TM6 not only shifted the activation curve but also changed its slope. A reduction in the mechanical sensitivity of the transduction channel, as represented by the slope of the activation curve, suggests that the force applied by the tip link is less efficiently coupled to the relative energies of the open and closed states. According to the classical gating-spring model ^48^, mechanical force is coupled to the channel through an elastic element known as the gating spring, which has not been molecularly identified. In this model, the slope factor Z of the activation curve (also known as the single channel gating force) is defined as:

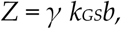

where *γ* is the geometric gain between bundle deflection and gating spring extension; *k*_*GS*_ is the gating spring stiffness; and *b* is the swing of the gate. Compared to mechanically tight hair bundles such as those from bullfrog, bundles in mammalian cochlea are difficult to stimulate with uniform deflection^49, 50^. Uneven contact of hair cells with the stimulus probe results in underestimation of the Z value, by as much as 85% for IHCs^49, 50^. Still, it is possible to compare the Z values from hair cells expressing mutated TMC1 channels and to determine if Z has been altered by the mutations.

In the simple biophysical model, a change in Z may be due to a change in the gating spring stiffness or in the swing of the gate. It is difficult to imagine how the mutation of a single residue in TMC1 could reduce the stiffness of the gating spring; whatever it is, the gating spring must be able to stretch many tens of nanometers and the structure of TMC1 is not conducive to such a huge rearrangement. From biophysical studies of bullfrog hair bundles, the gating swing has been estimated as 4 nm or more^51, 52^, a distance much larger than the ∼0.5 nm movement suggested by structural modeling that may close the outer part of the pore (Fig. 1D). Thus it is likely that a large movement of one part of the channel propagates to the gate region to open the pore itself. Reduction in Z by mutation could also come about in how efficiently the force produced by the large gating movement changes channel open probability.

We have explored the possibility that D528 is coupled to the mechanical stimulus via salt-bridge interactions with nearby positive charges with the protein itself. However, we find that mutations of R601 led to rightward shifts in the activation curves without a change in slope. This suggested that the proposed salt bridge between D528 and R601 may be important for stabilizing the closed channel conformation but doesn’t seem to play a role in the way the gate is coupled to the mechanical stimulus.

A number of proteins together form the mechanotransduction complex^11^. Previous studies have shown that TMC1 is the major pore-forming component of the complex. TMC1 might also be the force-sensing component of the complex, but it is also possible that tip-link tension is conveyed to a distinct force-sensing protein within the complex, and that a conformational change of that protein then promotes opening of the TMC1 pore ^53^. The block of gating compliance by pore-blocking compounds suggests that the pore is intimately associated with the conformational change produced by tension ^51, 54^. By changing the slope factor Z with single residue mutations in TM4 and TM6, this study specifically implicates TMC1 as part of the molecular sensor of tension. Based on this result, on ab inito structural modeling, and on gating movements in related channels, we propose also that the gating transition in TMC1 involves a separation of TM4 from TM6, to open the permeation pathway.

## Methods

### Structure predictions

We compared our previously reported homology model for TMC1 ^6^ with the newly generated and more refined models by the transform-restrained trRosetta^18^, RoseTTAFold^19^ and AlphaFold2^21^. The template-free predictions with AlphaFold2 were performed in a Colab notebook setting (Google Research). To obtain multiple conformations, we ran the network multiple times with different seeds. We typically investigated the top 10 of the ranked structures. Structures were aligned in and visualized by PyMOL (Version 2.4, Schrödinger). We aligned and clustered the structures using the program Chimera^2^. The permeation pathways were modeled by the software HOLE ^28^.

### Sequence alignment and evolutionary coupling analysis

Using EVcouplings software^55^ (beta version of v0.1.1), which uses the Markov model-based sequence-search tool JackHMMER^56^, we obtained a multiple sequence alignment of 5380 sequences. Alignments were built from the Uniref100^57^ dataset downloaded on July 24, 2020 using a maximum of 5 JackHMMER iterations. For further processing in the EVcouplings pipeline, we imposed a fragment requirement for each sequence to align to at least 70% of the query length, and analysis was restricted to just residue positions that contained at least 70% non-gaps. We used the top 100 covarying residue pairs (evolutionary coupligns) for further analysis.

### AAV virus preparations

Mutations were introduced to specific locations in the mouse AAV-Tmc1ex1 vector, carrying a cytomegalovirus (CMV) promoter ^6^. We generated TMC1 constructs bearing the following amino acid substitutions: G518A, G539A, G539I, G539/A544G, D528A, D528N, D528E, 601A, R601H, R523H, G411A, R601F/D528N, A544G (GenScript, Piscataway NJ, USA). Constructs were screened with SmaI digest to check for inverted terminal repeat (ITR) integrity before packaging into AAV vectors. Viral vectors were packaged with AAV2 ITRs into the AAV9 capsid serotype PHP.B. Virus were generated and purified by the Viral Core at Boston Children’s Hospital with genomic titers (measured with hGH primers) between 1 ×10^13^ and 5×10^14^ gc/ml. Vectors were aliquoted and stored at -80°C.

### AAV virus injections

Neonatal *Tmc1*^Δ/Δ^/*Tmc2* ^Δ/Δ^ C57BL/6 mice^34^ typically born ∼ E20, were placed with CD1 foster mothers. All animals were housed and bred in the animal facility at Harvard Medical School. Male and female pups were randomly chosen for the study. Prior to injection, mice were anesthetized using hypothermia. A post-auricular incision was made near the left ear and a cotton ball was inserted to spread the tissue. For round window membrane (RWM) injections, an injection needle made from a 1.4 mm glass capillary and pulled to a ∼10 µm diameter tip was positioned just above the RWM to confirm that viral suspension was released when the injection was started. Once confirmed, the needle was inserted through the RWM and 1.2 µl was injected using a Nanoliter 2000 Injector (World Precision Instruments) at a rate of 65 nl/min. Following the procedure, the surgical incision was closed with sutures. The pups were then put on a 37°C heating board to recover and returned to their cages. Usually three animals were injected in a session, and we typically performed three injection sessions per mutation.

### Cochlear dissections

At P4 to P6, pups were euthanized by rapid decapitation, temporal bones were dissected and the membranous labyrinth was isolated under a dissection microscope. Reissner’s membrane was peeled back, and the tectorial membrane and stria vascularis were mechanically removed. Organs of Corti were excised and cultured in medium containing DMEM supplemented with 1% FBS, 10 mM HEPES and 0.05 mg/ml carbenicillin at 37°C in 5% CO_2_. Organ of Corti cultures were pinned flat beneath a pair of thin glass fibers glued at one end with Sylgard to an 18-mm round glass coverslip. The tissue was placed in culture for 3-7 days before electrophysiological studies.

### FM1-43 uptake assays

Coverslips with adherent cochlear cultures were placed on a glass-bottomed chamber. The culture media was washed away with Leibovitz’s L15 medium three times and then incubated with 2 µM FM1-43 in L-15 for 1 minute followed by incubation with 2 mM SCAS in L-15 for 3 minutes and washed with L-15 two additional times before imaging.

### Confocal imaging

Imaging was performed on an Olympus FV1000 confocal microscope. FM1-43 fluorescence (excitation at 488 nm with ∼5-12 % intensity) was measured on mounted cultures with a 60X (1.1-NA) water-immersion objective.

### Electrophysiology

Transduction currents were recorded from IHCs using a Nikon Eclipse FNI microscope with 60X LWI objective and DIC optics and an Axopatch 200B patch clamp with a Digidata 1440 digitizer controlled by pCLAMP 10 software (Molecular Devices). The whole-cell voltage-clamp configuration was used for recordings. Currents were filtered at 5 Hz with a low-pass eight-pole Bessel filter. For hair-bundle stimulation, custom glass probes were made and polished to a diameter of ∼4 µm to match the shape of the inner hair cell bundles. The probe was attached to the probe holder with wax and shielded with grounded aluminum foil. The holder was moved by a piezo stack (Physik Instruments) driven by a custom high voltage piezo driver amplifier. Bundles were displaced for 80 ms with 15 step displacements from -175 nm to 1050 nm at 88 nm increments. For recordings, 1.5 mm OD R-6 (8350) glass pipettes were pulled with a Narishige PC-10 puller and coated with pre-warmed wax before use. These patch pipettes were filled with an internal solution containing (in mM): 137 CsCl, 5 EGTA, 10 HEPES, 2.5 Na_2_-ATP, 0.1 CaCl_2_ and 3.5 MgCl_2_, and adjusted to pH 7.4 with CsOH, ∼290 mmol/kg. The tissues were bathed in external solution containing (in mM): 137 NaCl, 5.8 KCl, 0.7 NaH_2_PO_4_, 10 HEPES, 1.3 CaCl_2_, 0.9 MgCl_2_, 5.6 glucose, vitamins and essential amino acids, adjusted to pH 7.4 with NaOH, ∼310 mmol/kg. Cells were held at a -80 mV potential and a separate pipet flowed extracellular solution onto their apical surfaces.

### Electrophysiology analysis

Data were analyzed with Clampfit (Molecular Devices) and ORIGIN (OriginLab). The average maximum current amplitudes as a function of probe displacement were plotted for each hair cell and fitted with a Boltzmann equation using ORIGIN as follows:

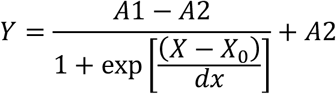

The data were then normalized for each cell, such that A1 and A2 range from 0 to 1 to allow comparison across cells. Z values were derived from the relationship 1/dx=Z/*k*_*B*_*T*. For each TMC1 variant, the average X_0_ and Z values were calculated as the mean ± SEM of all cells. The number of cells (n) is indicated in the text and figures. Student’s t-test was used to compare the means, and p values < 0.05 are marked as significant. For mutations with no significant change in slope, the corresponding change in intrinsic energy difference was estimated as Δ*X*_0_/*dx* in units of k_B_T. For non-stationary noise analysis, the variance of the responses was plotted against the mean for each cell. As before, we averaged the data within 1 pA intervals ^6^ and fitted with a parabola to derive the relevant values:

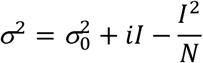

where σ2 is the variance, i is the single-channel current, I is the whole-cell current and N is the number of channels^39^. Cells were included in the analysis only if the plot of σ^2^ vs. I yielded a parabolic dataset that was fit well by the equation above. Statistics are based on at least three cells for each condition.

## Author contributions

N.A., B.P, K.D.K. and D.P.C designed the experiments. B.P. acquired electrophysiology data. K.D.K. assembled instrumentation and performed FM1-43 assays and confocal imaging. P.T. performed cochlear dissections. Y.L. bred and maintained mice and performed cochlear injections. K.P.B. and D.S.M mapped evolutionarily coupled residue pairs onto structures. N.A., K.D.K., B.P., and D.P.C wrote the manuscript.

## Acknowledgements

We thank Drs. Ivan Anishchenko, Artur Indzhykulian, Chuck Phillips, Marcos Sotomayor and Jeffrey Lottheimer for helpful discussions; Yiming Zhang for producing AAV vectors; Bruce Derfler for extensive assistance with molecular preparations; and Drs. Andrew Griffith and Jeffrey Holt for providing *Tmc1*^Δ/Δ^/*Tmc2* ^Δ/Δ^ mice. This work was supported by the NIH (R21 DC018631 to NA and R01 DC000304 to DPC).

