## Supplementary Information for "Mechanical gating of the auditory transduction channel TMC1 involves the fourth and sixth transmembrane helices"

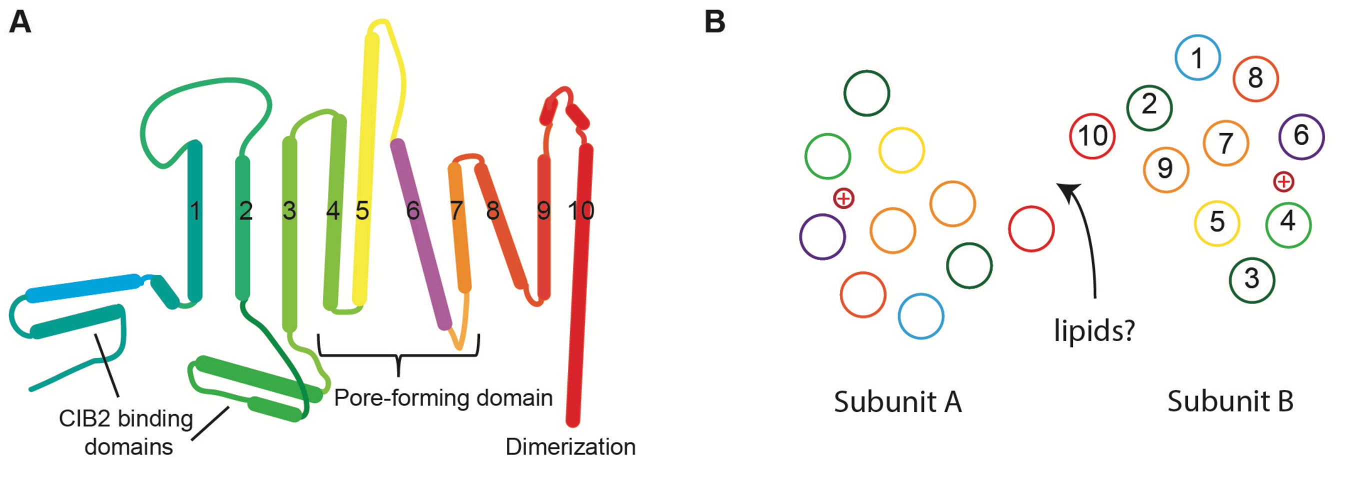


**Supplementary Fig. 1:** A) Schematic representation of the 10-TM topology of TMC1. B) Schematic for the predicted arrangement of the helices in the dimeric structural models. A top to down view of the dimeric channel is shown with each TM helix represented as a circle and colored as in panel A. Subunit B shows TM helix numbers. The approximate position of the pore cavity is marked with a + sign in each subunit.

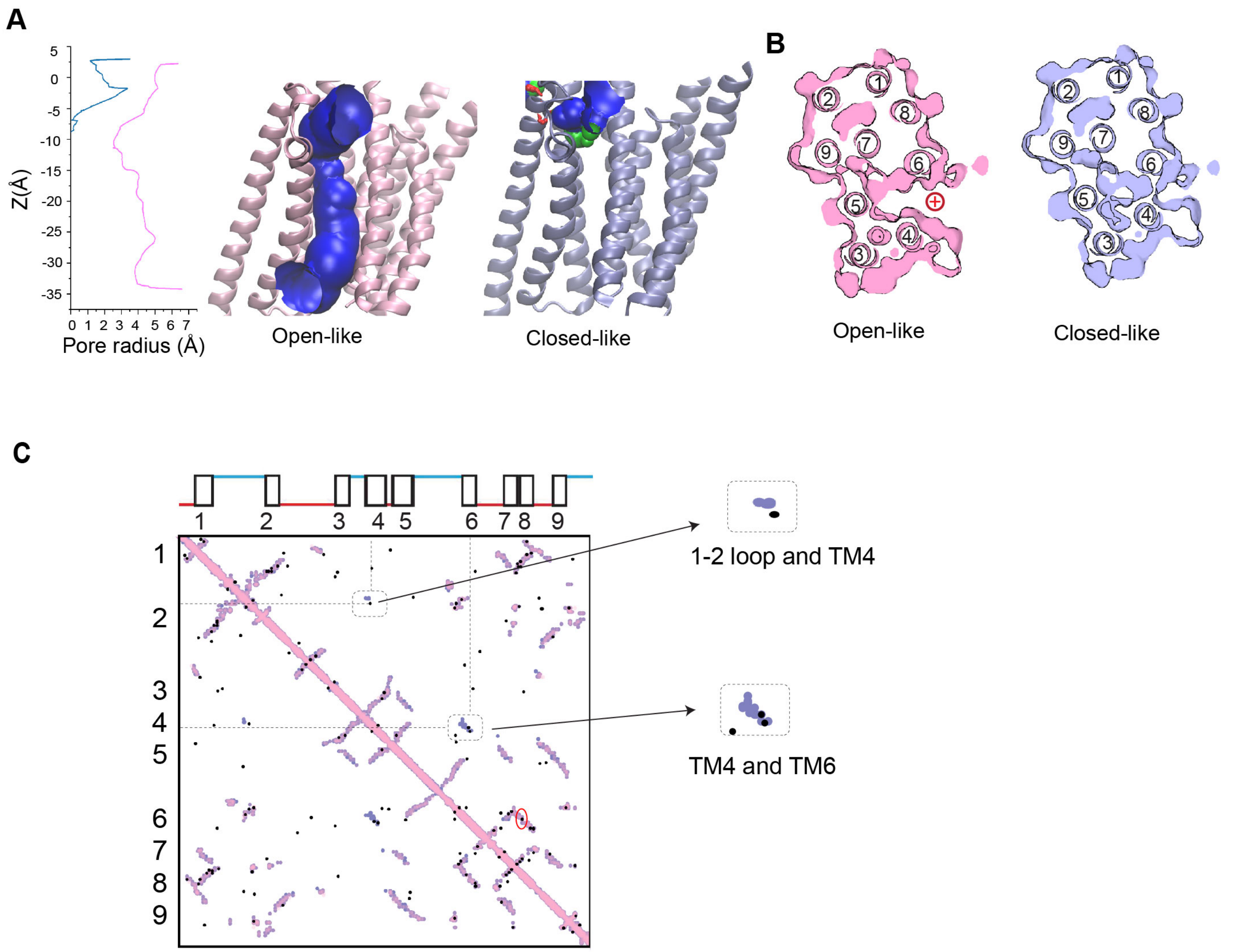


**Supplementary Fig. 2. A)** The permeation pathway through the predicted open-like (pink) and closed-like (blue) structures. The estimated pore radius is plotted against the pore length Z, from outside to inside. The approximate position of the constriction site in the open-like structure, about 10 Å in, corresponds to approximately N404 on TM4 and L524 on TM6. Loops and termini (including residues 1-174, 212-266, 294-351, 466-511, 545-566, 617-625, 655-696, 725-756) are not shown for simplicity. **B)** A thin cross section of the protein parallel to the membrane at the Z-level of approximately N404 on TM4 and L524 on TM6 for the open-like and closed-like structural predictions (pink and blue, respectively**)**. **C)** Contact maps for residues 165-680 in TMC1 predicted by EVcoupling analysis (black dots) are mapped onto contact maps from open-like and closed-like predicted structures (pink and blue, respectively). Evolutionarily coupled (EC) pairs that are found in proximity only in the closed state are indicated to the right: they connect TM4 to TM6 (notably S408-T531 and F413-Y533), and the TM1-2 loop to TM4 (Q245 on TM1-2 loop and M403 on TM4). The transmembrane domains corresponding to the residues along the axes are numbered 1-9. The EVcoupling analysis suggests that these residue pairs are likely to interact; the AlphaFold2 structures also predict interaction but only in the closed-like conformation. EC 528-601 between TM6 and TM8, which is the second strongest coupling out of all possible analyzed pairs, is circled in red.

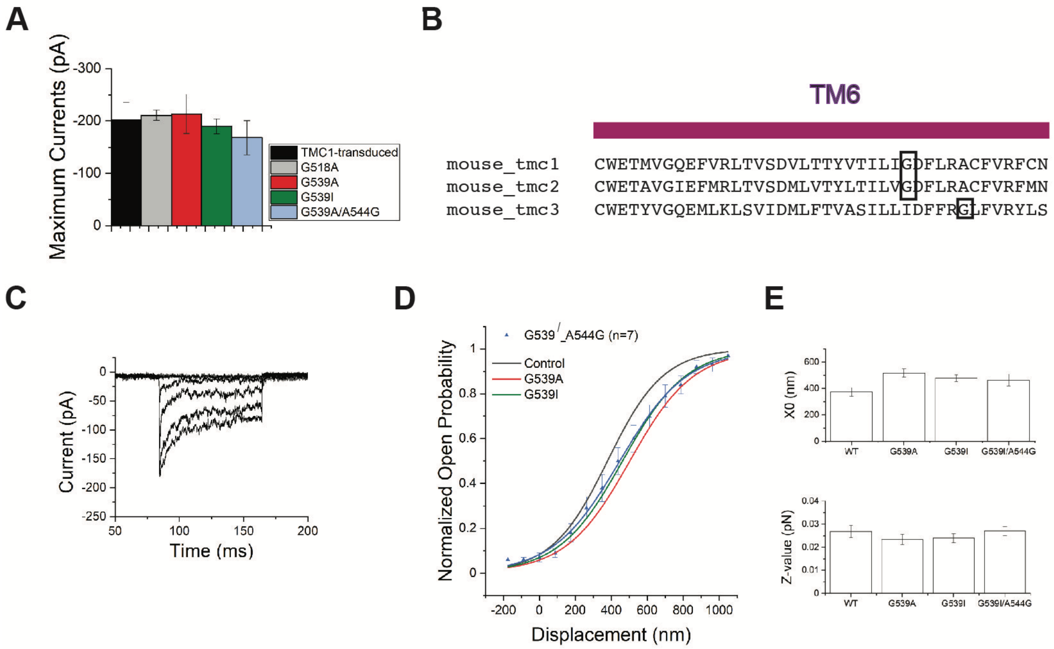


**Supplementary Fig. 3. A**) The average maximum transduction currents for hair cells virally expressing WT-TMC1, TMC1-G518A, TMC1-G539A, TMC1-G539I and TMC1-G539A/A544G. Amplitudes are all similar. **B**) Sequence alignment of the mouse TM6 helix in orthologs TMC1, 2 and 3. TMC1 and TMC2 have glycine at position 539; TMC3 does not but has a glycine at position 544. **C**) Representative currents from hair cells expressing TMC1-G539I/A544G. **D**) Normalized open probability curve for G539I/A544G compared to the WT control, G539A and G539I. **E**) Fitting parameters for the four TMC variants. None are significantly different (p =0.1, 0.4 and 0.8 for $X_0$, and 0.7, 0.6 and 0.7 for Z, relative to WT).

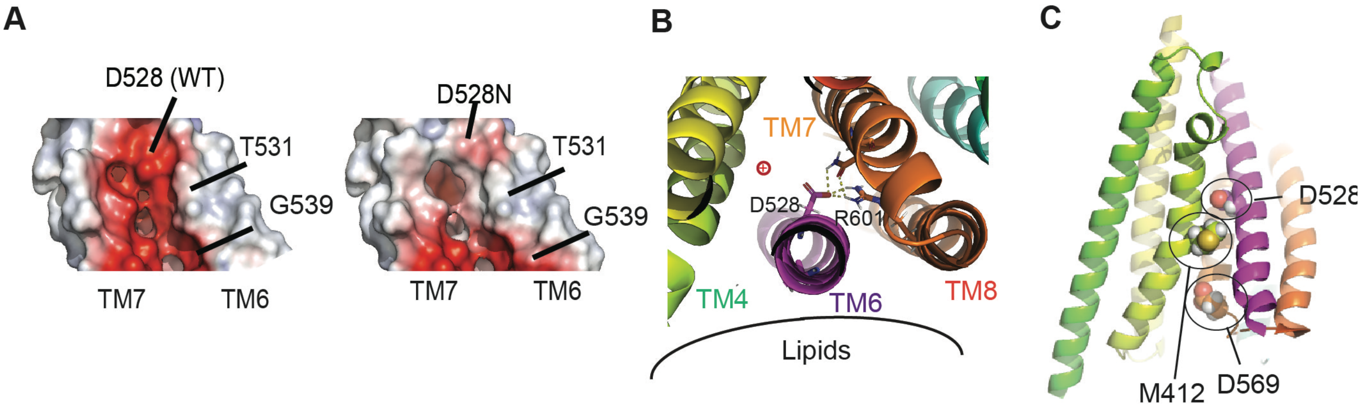


**Supplementary Fig. 4. A)** Electrostatic surface charge distribution of pore-lining TM6 and TM7 helices, viewed from inside the pore. TM6 contacts the lipid, TM7 is within the protein core. WT-TMC1 and TMC1-D528N structures were generated with AlphaFold2. The D528N mutation substantially reduces the negative charge that might enhance passage of cations. **B)** Extracellular view of the selected residues D528 and R601. The approximate locations of the permeation pathway (**+**) and the lipid bilayer (black curve) are indicated. In this conformation, the side chain of D528 forms a salt bridge with the side chain of R601. **C)** The position of deafness mutations M412 and D569 are indicated. In the closed-like conformation, these sites would be inaccessible to extracellular solution (Supplementary Fig. 2A), consistent with their limited accessibility in the closed state as assessed with cysteine-modifying reagents (Pan et al., 2018).

**A**

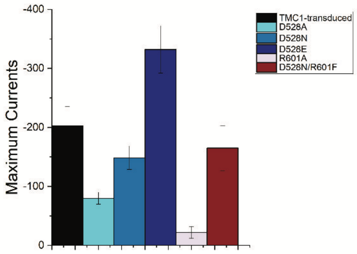


**Supplementary Fig. 5.** Average maximum transduction currents for hair cells virally expressing WT-TMC1, TMC1-D528A, TMC1-D528N, TMC1-D528E, TMC1-R601A and TMC1-D528N/R601F. There is variability among these variants in whole cell currents among. Variability is expected because most of the mutations are likely to alter the charge distribution within the permeation pathway.

**Supplementary Table 1:** TMC1 Mutants Packaged in AAV9-PHP.B

| Mutation | Titer (vg/ml) | Location |
|---|---|---|
| G518A | 1.19E+13 | TM6 |
| G411A | 3.04E+14 | TM4 |
| G539A | 7.65E+13 | TM6 |
| G539I/A543G | 8.64E+13 | TM6 |
| A543G* | 2.70E+14 | TM6 |
| G539I | 1.76E+14 | TM6 |
| D528A | 4.83E+13 | TM6 |
| D528N | 3.69E+13 | TM6 |
| D528E | 1.48E+14 | TM6 |
| R601A | 6.36E+13 | TM8 |
| R601H | 4.50E+14 | TM8 |
| R601F/D528N | 2.17E+14 | TM8/6 |
| R523H | 1.66E+14 | TM6 |
| WT-TMC1-HA | 3.12E+13 | N/A |

*** Data not shown.**

**Supplementary Table 2:** Occurrence of the indicated amino acids at the positions corresponding to 518, 539 and 544. The amino acids found in the mouse TMC1 sequence are underlined. The frequencies represent those occurring in a multiple sequence alignment of 5,050 TMC sequences (see Methods).

| | G518 | G539 | A544 |
|---|---|---|---|
| A | | | **<u>27%</u>** |
| G | **<u>98%</u>** | **<u>50%</u>** | 11% |
| R | | | 34% |
| K | | | 19% |
| I/V | | 41% | |

**Supplementary Table 3:** Frequency of finding the indicated amino acids at the positions corresponding to 523/528/601 within the mouse TMC1 sequence. The frequencies represent those occurring in a multiple sequence alignment of 5,050 TMC sequences (see Methods).

| 523/528/601 | |
|---|---|
| K/D/K | 30% |
| R/D/K | 20% |
| R/D/R | **17%** |
| K/D/R | 7% |
| K/N/F | 6% |
| other | 20% |

**Supplementary Video 1:** A possible gating trajectory, if the two groups of AlphaFold2 structures represent closed and open states of TMC1.
